# Impact of Antibiotics on the Genomic Expression of *Pseudomonas aeruginosa* in the East African Community: A Systematic Review

**DOI:** 10.1101/2024.12.23.630126

**Authors:** Comfort Danchal Vandu, Ilemobayo Victor Fasongbon, A. B. Agbaje, Chinyere Njideka Anyanwu, Makena Wusa, Emmanuel O. Ikuomola, Reuben Samson Dangana, Nancy B. Mitaki, Ibe Micheal Usman, Augustine Oviosu, Herbert Mbyemeire, Elizabeth Umorem, Shango Patience Emmanuel Jakheng, Musyoka Angela Mumbua, Solomon A Mbina, Esther Ugo Alum, Ibrahim Babangida Abubarkar, Swase Dominic Terkimbi, Siida Robert, Ezra Agwu, Patrick Maduabuchi Aja

## Abstract

Antimicrobial resistance (AMR) presents a significant health problem globally with the majority of the burden coming from lower-middle-income countries. AMR surveillance under a One Health paradigm is critical for determining the relationships between clinical, animal, and environmental AMR levels. Allowing for a thorough knowledge of the interconnected variables contributing to resistance, which enables the development of effective solutions.

This systematic review was conducted to determine the impact of antibiotics on the gene expression of *Pseudomonas spp.* In the East African Community. A comprehensive literature search was conducted across Web of Science, Scopus, and PubMed databases yielding 284 articles with 11 meeting the inclusion criteria after screening.

We included the 11 studies from 5 East African Countries that are part of the East African Community, the results revealed a high prevalence of antimicrobial resistance in *Pseudomonas aeruginosa*, with resistance rates above 90% for most tested antibiotics, exception of Amikacin, which remained effective due to its limited use. Common resistance genes reported included carbapenem-resistant genes like *blaNDM-1* and *blaVIM*, the most common method used was disc diffusion method at (50%). The review also found high-risk clones, such as ST 244 and ST 357, that were associated with multidrug-resistant strains. Environmental isolates showed lower resistance rates (54%) than clinical pathogens (73%), indicating different selecting pressures. Majority of the studies were conducted in Kenya (30%) and Uganda (30%), indicating differences in research capabilities and healthcare facilities.

These findings highlight the critical need for more surveillance, effective antimicrobial stewardship programs, and additional research to prevent antibiotic resistance and guide public health initiatives in the region.

**KEY FINDINGS OF THE STUDY:** *Pseudomonas aeruginosa* isolates demonstrated substantial resistance to antibiotics, including cefepime, meropenem, levofloxacin, and ticarcillin-clavulanic acid as reported across various studies conducted in East Africa.

Amikacin was reported to be more effective in more than 90% of the studies reported across East Africa as a potential treatment choice for multidrug-resistant *Pseudomonas* infections in the region.

Carbapenem-resistant genes such as *blaNDM-1, blaVIM*, and *blaOXA-48* were found in a large number of clinical and environmental isolates.

High-risk clones, such as ST 244 and ST 357 were reported to demonstrate clonal spread of multidrug-resistant *Pseudomonas aeruginosa* across East African healthcare settings.

The disc diffusion method was the most popular antimicrobial susceptibility testing method (50%), owing to its low cost and simplicity.

DNA extraction and PCR were used in 30% of the studies whereas more advanced approaches such as whole genome sequencing were less popular due to resource constraints.

The majority of studies were undertaken in Kenya (30%) and Uganda (30%), with fewer studies in Tanzania and the Democratic Republic of the Congo (20%), demonstrating regional variations in research capacity and healthcare resources.

## INTRODUCTION

*Pseudomonas aeruginosa* is one of the most common Gram-negative bacterium that is well-known for its flexibility and plasticity in a variety of contexts, including natural habitats and therapeutic settings (Darwesh, & Matter, 2024). Due to its potential to produce serious infections and its exceptional resistance to antibiotics, its study has attracted a lot of attention globally in recent years Hafiz *et al.,* (2023). It has been reported that *P. aeruginosa* is an opportunistic pathogen that mostly affects persons with impaired immune systems. It is commonly linked to hospital-acquired diseases, such as bloodstream infections, pneumonia, and urinary tract infections (Salim, Mohan, Forgia & Busi., 2024). The numerous resistance mechanisms and environmental adaptability of this bacteria pose significant hurdles in clinical settings (Wood, *et al.,* 2023). Through a variety of strategies, such as overexpressing the efflux pump, changing target locations, and forming biofilms, the bacteria can modify its gene expression to acquire resistance (Zhang, *et al.,* 2024). Because of its intricate regulatory networks that manage gene expression, *P. aeruginosa* can withstand a variety of drugs and thrive in hostile environments (Mudgil, Khullar, Chadha, & Harjai *et al.,* 2024).

Exposure to antibiotics can significantly alter *P. aeruginosa* gene expression profiles (Huynh, *et al.,* 2024). Genes linked to virulence, resistance, and biofilm formation may express differently in response to sub-inhibitory antibiotic doses (Marmolejo, 2024); exposure to sub-lethal concentrations of antibiotics can induce the expression of efflux pumps such as *MexAB-OprM* and *MexXY*, which actively expel antibiotics from the bacterial cell, thereby reducing their efficacy (Morita *et al.,* 2020; Santos, *et. al.,* 2024). For example, it has been demonstrated that exposure to aminoglycosides, like tobramycin, increases the expression of the *arr* gene, which is involved in the production of biofilms (David, *et. al.,* 2024; Abdullah, & Younis, 2024).). The intricacy of antibiotic-induced gene expression regulation is demonstrated by the similar differential gene expression responses to colistin and tobramycin, which affect both the transcriptional and translational levels (Blevins *et al.,* 2018; Araújo, *et. al.,* 2024; Wang, Gu, Zhang, & Zhu, 2024). In addition, it has been demonstrated that antibiotics such as ceftazidime and ciprofloxacin activate genes involved in stress response, which improve bacterial survival in harsh environments (Fernando *et al.,* 2021; Almutairy, 2024). *P. aeruginosa*’s gene expression is also highly influenced by environmental factors (Cecil, & Yoder-Himes, 2024). A variety of stresses, including variations in pH, temperature, and the availability of nutrients, can cause adaptive reactions in bacteria that increase their ability to survive and spread disease (Cardoza, & Singh, 2024). For instance, in response to external stimuli, the expression of genes involved in quorum sensing and biofilm formation is frequently elevated, which promotes the onset and persistence of infections (Erkihun, *et. al.,* 2024). Disinfectants and heavy metals are two examples of pollutants that can cause stress reactions and encourage horizontal gene transfer, which can help resistance genes proliferate (Ni, Zhang, Cai, Xiang, & Zhu, 2024). Research has indicated that prevalent disinfectants, quaternary ammonium compounds (QACs), can alter the composition of membranes and enhance the expression of genes related to resistance (Kim *et al.,* 2018; Lu, Mahony, Arnold, Marshall, & McNamara, 2024). According to Sauer *et al*. (2019) and Ungor, & Apidianakis, (2024) virulence factors can also be expressed in response to environmental factors including oxygen levels and nutrition availability, which can alter *P. aeruginosa*’s pathogenicity.

Antibiotic exposure and environmental factors can work in concert to improve *P. aeruginosa*’s capacity for adaptation (Ambreetha, Zincke, Balachandar, & Mathee, 2024; Jin, 2024). For instance, because antibiotic resistance genes frequently co-occur on the same genetic material that is mobile, the presence of heavy metals in the environment might co-select for these genes (Murray, *et. al.,* 2024; Goff, *et. al.,* 2024). According to Martínez *et al.,* (2020), the co-selection mechanism makes controlling *P. aeruginosa* infections more difficult by increasing the issue of antibiotic resistance. Furthermore, antibiotic pressure and environmental stimuli both affect biofilm development, a critical component of persistent infections that results in an increasing tolerance to antimicrobial drugs (Flemming *et al.,* 2019).

Significant clinical implications result from the interplay of *P. aeruginosa* gene expression and environmental variables in conjunction with antibiotic treatment (Yang, *et. al.,* 2024; Sendra, *et. al.,* 2024). Treatment plans may become more difficult to implement as a result of the adaptive reactions that environmental stresses and antibiotics can cause (Yang, Li, Yao, & Li, 2024; Kleinman, 2024; Almutairy, 2024). Subinhibitory antibiotic doses, for example, can encourage the production of biofilms, which improves the bacterium’s resistance to antimicrobial drugs and the human immune response (Guo, *et. al.,* 2024; Yousefi, *et. al.,* 2024). To effectively design therapeutic approaches, a thorough understanding of the molecular mechanisms underlying these adaptive responses is crucial due to their dual impact (Akbari, 2024).

## MATERIALS AND METHODS

### Literature Search

A comprehensive literature search was systematically conducted across the Web of Science (WoS), Scopus, and PubMed databases on May 23, 2024. The search employed the following terms: “antibiotics OR antibacterial,” “environment OR habitat,” “genomic OR DNA OR gene,” “pseudomonas,” and “east Africa.” Boolean operators (AND/OR/NOT), alternative terms, and various delimiters such as quotation marks, parentheses, wildcards, and asterisks (*) were utilized to create the search strategy outlined in **Table 1**, as reported by Fasogbon *et al*. (2022, 2023, 2024). The search was restricted to peer-reviewed articles published in English. The paper selection process adhered to the Preferred Reporting Items for Systematic Reviews and Meta-Analyses (PRISMA) 2020 guidelines (Page *et al.,* 2021; Mustapha *et al.,* 2022) and followed the inclusion and exclusion criteria presented in **Table 2**.

**Table 1.** Search Strategies

| Database | Search strategy |
| --- | --- |
| Scopus | ( antibiotic* OR antibacteria* OR clinical OR environment* OR surrounding* OR habitat* ) AND ( genom* OR dna OR "deoxiribonucleic acid" OR gene* ) AND ( pseudomona* ) AND ( congo OR drc OR somalia OR burundi OR kenya OR rwanda OR "south sudan" OR uganda OR tanzania OR "East Africa Community" ) |
| WoS | ( antibiotic* OR antibacteria* OR clinical OR environment* OR surrounding* OR habitat* ) AND ( genom* OR dna OR "deoxiribonucleic acid" OR gene* ) AND ( pseudomona* ) AND ( congo OR drc OR somalia OR burundi OR kenya OR rwanda OR "south sudan" OR uganda OR tanzania OR "East Africa Community" ) |
| PubMed | ((antibiotic*[title/abstract]) or (antibacteria*[title/abstract]) or clinical[title/abstract]) or (environment*[title/abstract]) or (surrounding*[title/abstract]) or (habitat*[title/abstract])) and ((genom*[title/abstract]) or (dna[title/abstract]) or (deoxiribonucleic acid[title/abstract])) and (pseudomona*[title/abstract]) and (((congo[title/abstract]) or (drc[title/abstract]) or (somalia[title/abstract]) or (burundi[title/abstract]) or (kenya[title/abstract]) or (rwanda[title/abstract]) or (south sudan[title/abstract]) or (uganda[title/abstract]) or (tanzania[title/abstract]) or (East Africa Community[title/abstract])) NOT (review[Publication Type])) |

**Table 2.** Inclusion and Exclusion Criteria

| Inclusion Criteria | Exclusion Criteria |
| --- | --- |
| Studies involving the expression of any virulence or antibiotic-resistant gene in any *Pseudomonas spp.*<br>Studies involving the impact of antibiotics or environment.<br>Studies conducted in East Africa Community.<br>Peer-reviewed articles published in English.<br>Studies involving the expression of any virulence or antibiotic-resistant gene in any *Pseudomonas spp.* | Studies not focusing on gene expression.<br>Studies that did not describe the impact of antibiotics or environment on gene expression.<br>Studies that are not on *Pseudomonas spp* were excluded.<br>Non-English publications.<br>Studies conducted elsewhere in East Africa Community. |

## RESULTS

The Web of Science database search produced 93 articles, Scopus yielded 180, and PubMed identified 11, bringing the total to 284 articles across the three databases. The search results from each database were then exported and imported into Rayyan, a platform specifically designed for systematic review processes, where they were screened based on the inclusion and exclusion criteria (Johnson & Phillips, 2018). On the platform, 70 duplicate articles were removed. The remaining 214 records were initially screened by evaluating their titles and abstracts, which led to the exclusion of 116 articles. Following a more detailed full-text review, an additional 87 articles were excluded for not meeting the inclusion criteria. In the end, 11 articles were included in the study after passing the eligibility criteria and quality assessment as presented in Figure 1.

**Figure 1:**
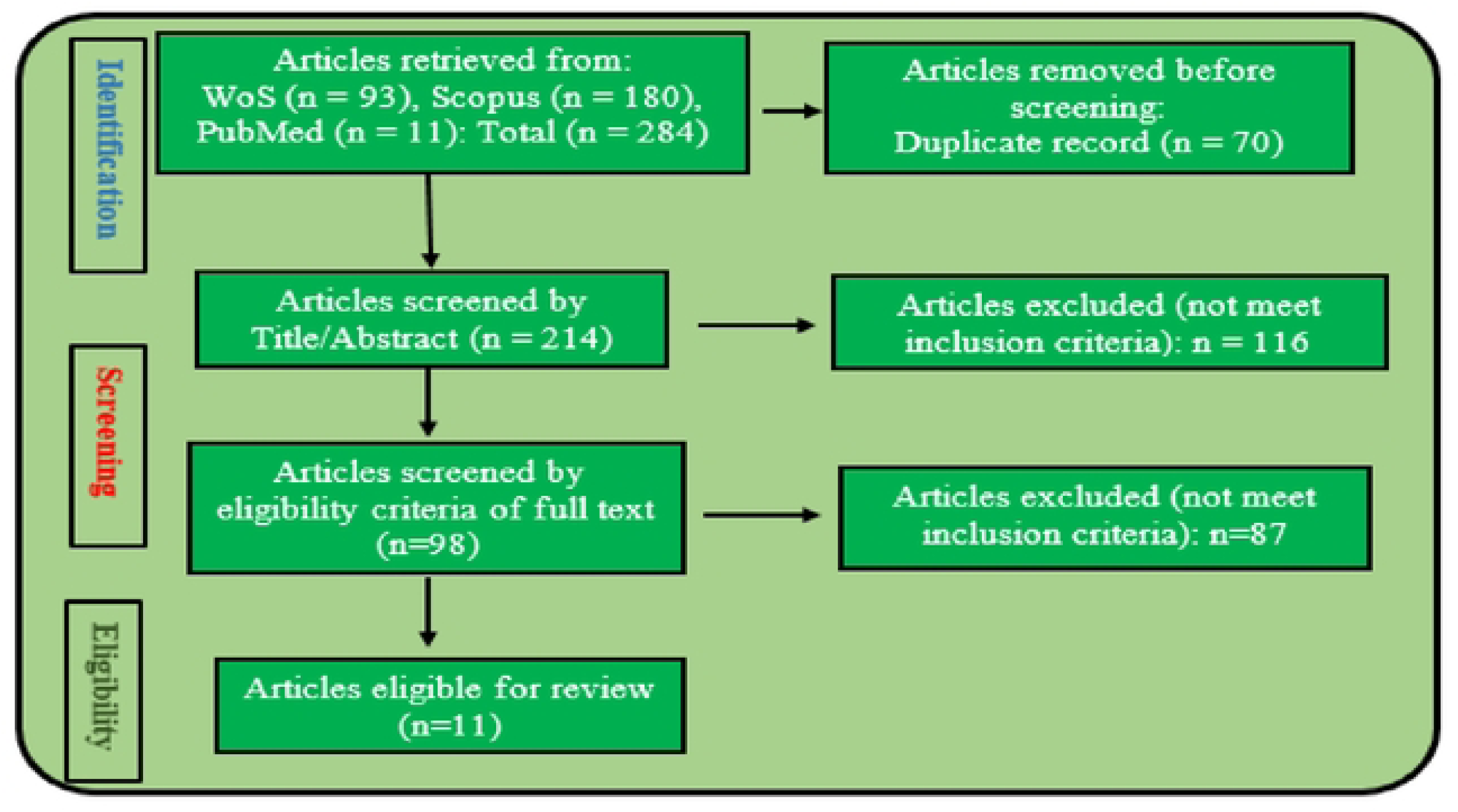
The PRISMA flow chart

Table 5, indicates that the highest number of studies were done in Kenya (4, 40%), followed by Uganda (3, 30%), while Tanzania and the Democratic Republic of Congo have (2, 20%), there were no eligible studies from Burundi, Rwanda, Somalia and South Sudan. These results indicate that there are only a few studies (not up to 50%) done on the impact of antibiotics or the environment, on the gene expression of *Pseudomonas spp* in the East African communities.

Figure 4 shows the methods used in the isolation and identification of *Pseudomonas spp* from the various samples in the studies, it shows that 70% of the studies used the traditional bacteria culture and biomedical tests to identify the organism, while only 10% used other methods as the Bactec automated system, bacterial culture for lipase activity (and under UV light), and the 16S rRNA sequencing for pathogen identification. The popular disc diffusion method of conducting antimicrobial sensitivity testing is used by most of the studies (50%) to identify the resistant Pseudomonas species, (20%) used Vitek and the others (10%) used Phoenix automated AST panel, broth microdilution, and agar diffusion methods each.

The most common antibiotic used to determine the resistance profile of the *Pseudomonas spp* is the Imipenem (90%), Amikacin, Ciprofloxacin, Cefepime, and Ceftazidime were used in (70%) of the studies; Ceftriaxone, Piperacillin-Tazobactam, and Gentamycin were used in (50%) of the Studies; Cefotaxime in (30%), while all the other antibiotics (Erythromycin, Clindamycin, Cefoperazone + Sulbactam, Cefpodoxime, Cefixime, Ampicillin, Amoxicillin-Clavulanic acid, Cefuroxime, Ticarcillin-Clavulanic acid, Levofloxacillin, Tetracycline, Tigecycline, Minocycline, Aztreonam, Cloxacillin, Zincsulphate, and Colistin) were each used in (10%) of the studies (Figure 2). For the methods of genomic characterization, (30%) of the studies used DNA extraction and PCR, (15%) used Whole Genome Sequencing and the Modified Hodge Test for Carbapenemase activity; and (5%) of the studies used only PCR, Quantitative PCR, and Imipenem-EDTA double disk synergy test for metallo-B lactamase activity (Figure 5).

**Figure 2:**
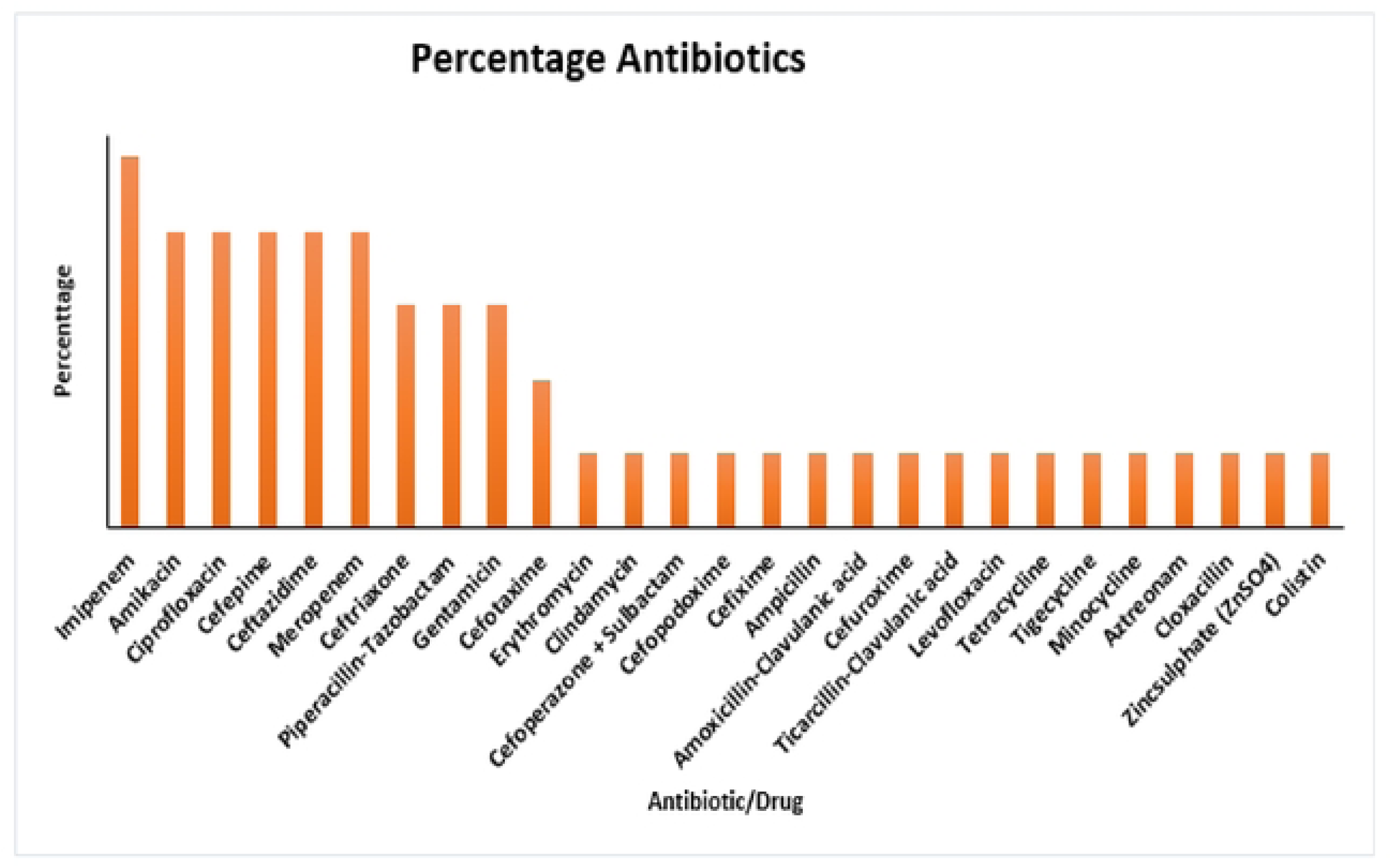
Percentage of Antibiotics reported across various Studies

From Table 4, the different strains of *Pseudomonas aeruginosa* isolated are, the global high-risk strains (ST 244 and ST 357;56). From the clinical samples; the high-risk strains (ST 357, ST 654), Local High-risk strains (ST 2025, ST 455 & ST 233), Carbapenem-resistant *P. aeruginosa* (ST 316, ST 357, ST 654 & ST 1203;13) were identified. Also, the novel multidrug-resistant clone with a virulence gene (ST 3674) was identified. While in all studies where other species of *Pseudomonas (P. pseudoalcaligens)* were isolated, no further characterization was done to identify the strains.

**Table 3:**
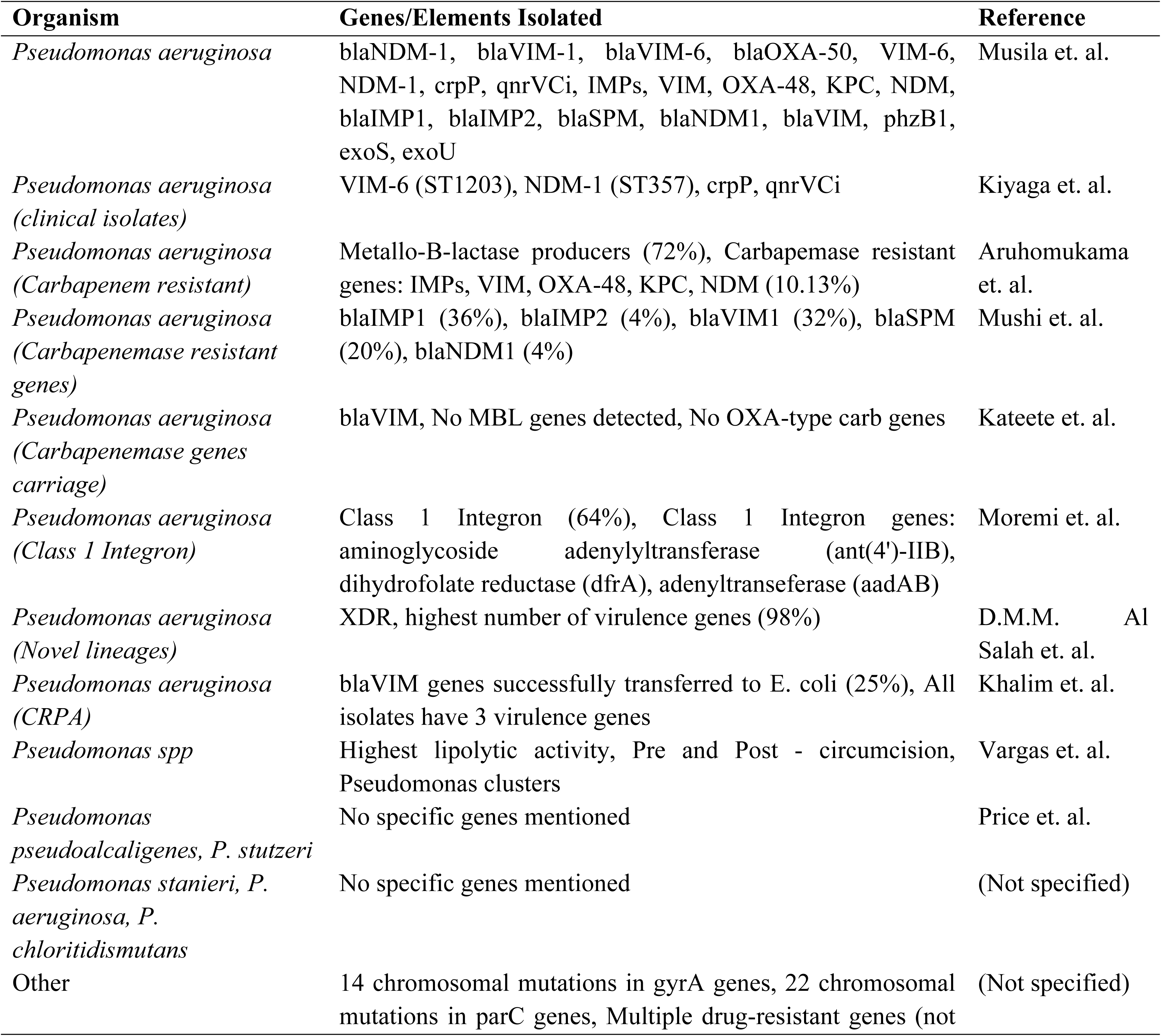
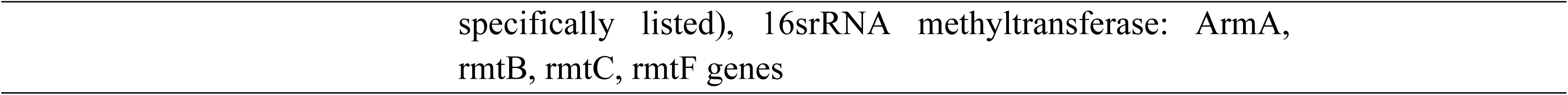
*Pseudomonas* Resistance and Virulence Genes Detected

**Table 4:** *Pseudomonas* Strains Sequenced

| Organism | Strain | Reference |
| --- | --- | --- |
| *Pseudomonas aeruginosa* | Global High-risk clones: ST 244 & ST 357; 56 *P.aeruginosa*, clinical isolates; High Risk STs: 357, 654; Local High-risk clones: ST 2025, ST 455 & ST 233; Carbapenem resistant *P. aeruginosa*; STs 316, 357, 654 & 1203; 13 *Pseudomonas aeruginosa* isolated; Novel MDR clone with virulence genes: ST 3674 | Kiyaga et. al.,<br>Musila et.al.,<br>Aruhomukama et.al. |
| *Pseudomonas aeruginosa* | NA | Moremi et.al.,<br>Khalim et. al.,<br>Irenge et.al., Vargas<br>et.al., Price et.al. |
| *Pseudomonas pseudoalcaligenes* | NA | D.M.M. Al Salah<br>et.al. |

**Table 5:** Countries and Number of Studies reported

| Country | Frequency | Percentage | Region |
| --- | --- | --- | --- |
| DRC | 2 | 20% | Kinshasa, South Kivu District |
| Kenya | 4 | 40% | Lake Bogoria of the Great Rift Valley, Nairobi (Agha Khan hospital), Six hospitals in Nairobi & Kisii |
| Tanzania | 2 | 20% | Northwestern, Northwestern region (Mwanza Hospital) |
| Uganda | 3 | 30% | Mbarara Referral Hospital, Mulago hospital Kampala |

## Discussion

This systematic review was conducted to identify the impact of antibiotics and the environment on the gene expression pattern of *Pseudomonas spp.* This integrates antimicrobial resistance patterns, genomic characterization, and the influence of environmental factors on bacterial behavior. *Pseudomonas aeruginosa* clinical isolates demonstrated a high level of resistance, with resistance observed in nearly all tested antibiotics except Amikacin in some studies (Kiyaga et al.,2022). Antibiotics that demonstrated resistance as reported across various studies include, Cefepime, Ticarcillin-Clavulanic acid, Piperacillin, Meropenem, Levofloxacin, Tetracycline, Tigecycline Minocycline according to Musila et al. (2021) and Kiyaga et al. (2022). The resistance reported cuts across multiple antibiotic classes, including cephalosporins (e.g., Ceftriaxone, Cefepime), carbapenems (e.g., Imipenem, Meropenem), and fluoroquinolones (e.g., Levofloxacin), with resistance varying across isolates and geographic locations as shown in **Figure 2**. The disc diffusion method was the most common method utilized, accounting for 50% of the studies. These could be due to its simplicity, cost-effectiveness, availability, and long-standing usage in clinical microbiology to provide reliable results easy to interpret with minimum requirement. More modern methods, such as the Vitek 2 system, were employed in 20% of research, while the Agar diffusion, Broth microdilution, and Phoenix automated AST panel methods were each used in 10% of the studies indicating resource availability differences as summarized in F**igure 3**.

**Figure 3:**
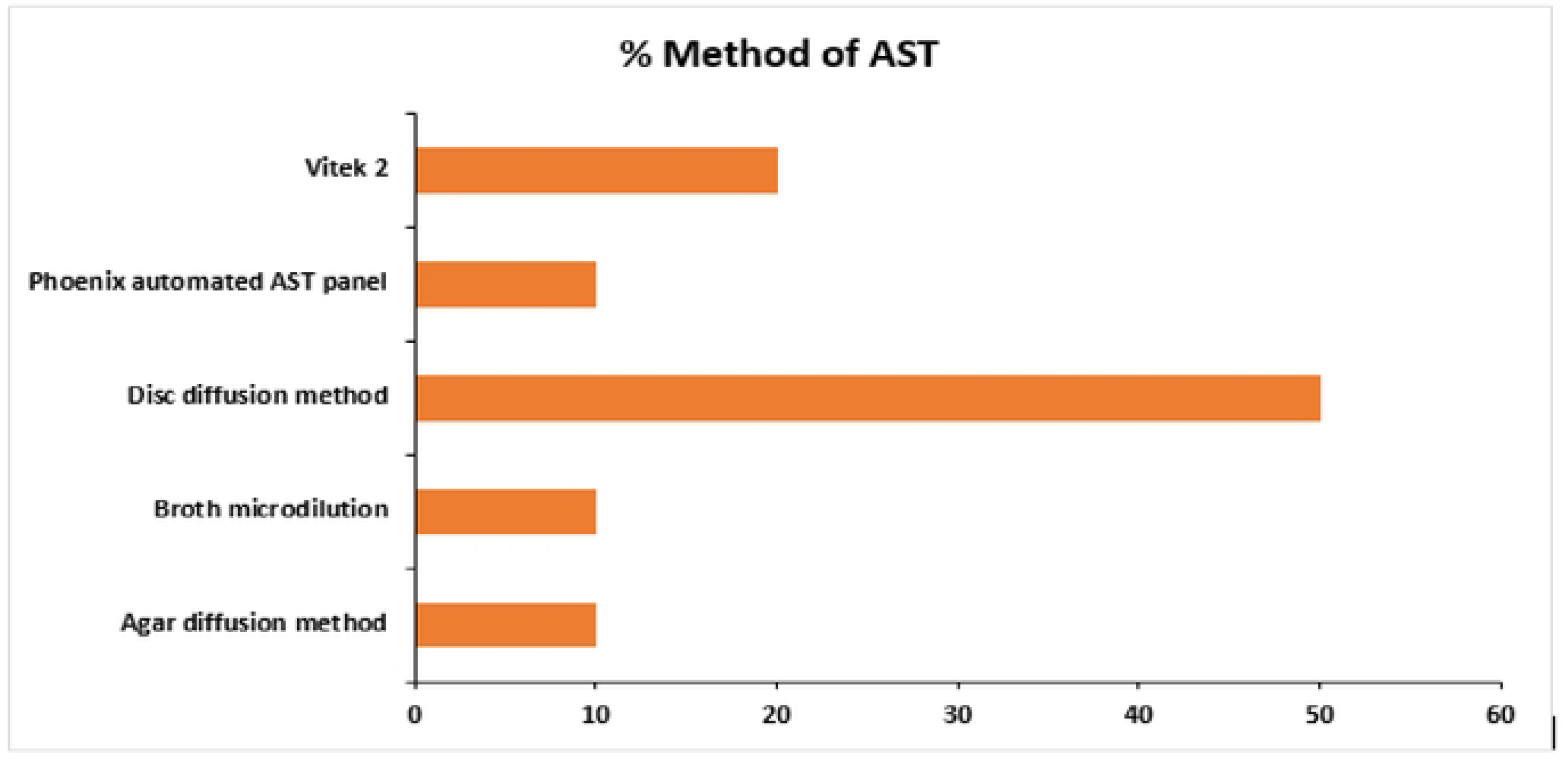
Methods of Antimicrobial Sensitivity Tests

**Figure 4:**
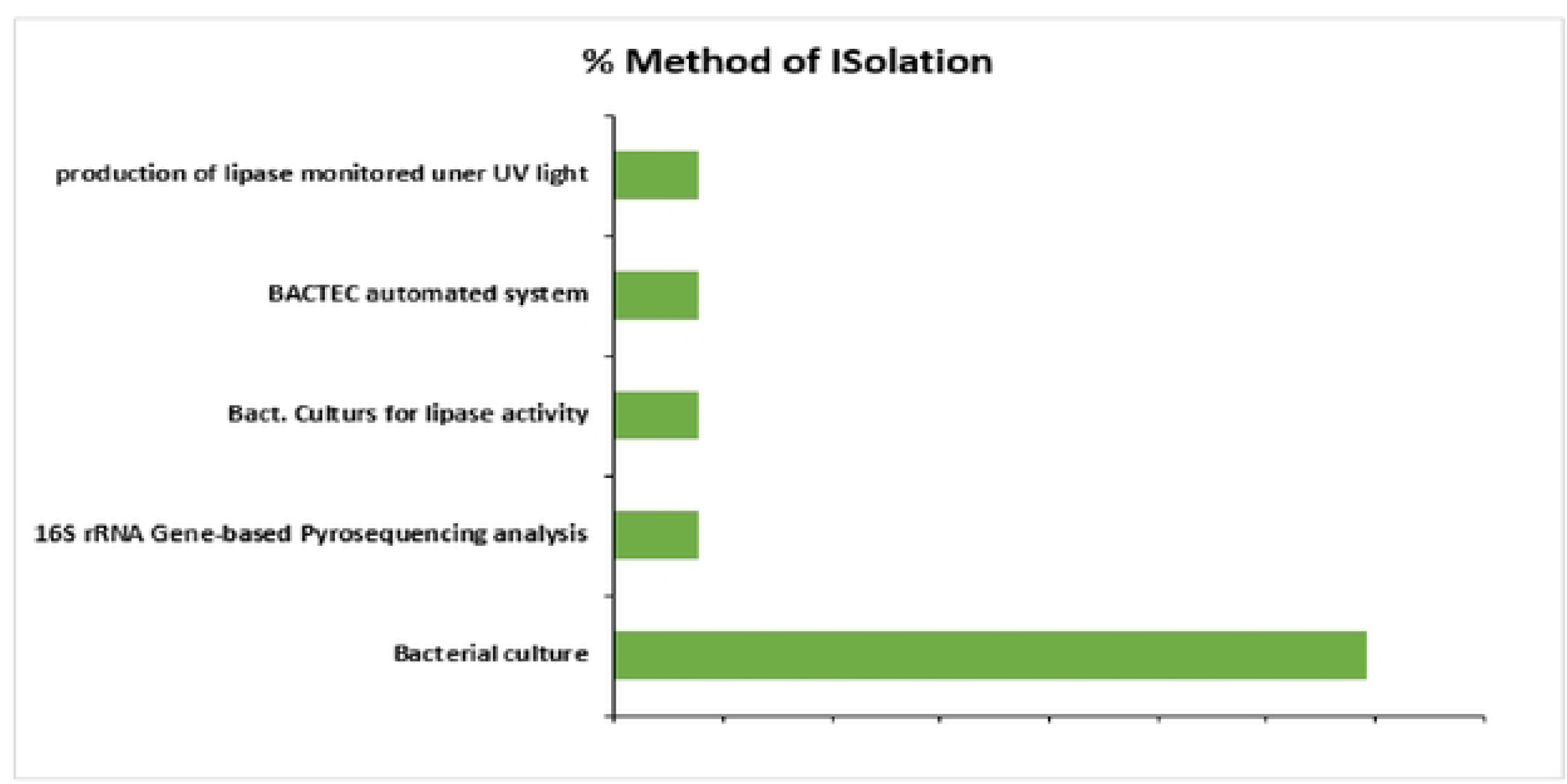
Methods used in the isolation and identification of *Pseudomonas :.pp* reported across various Studies

**Figure 5:**
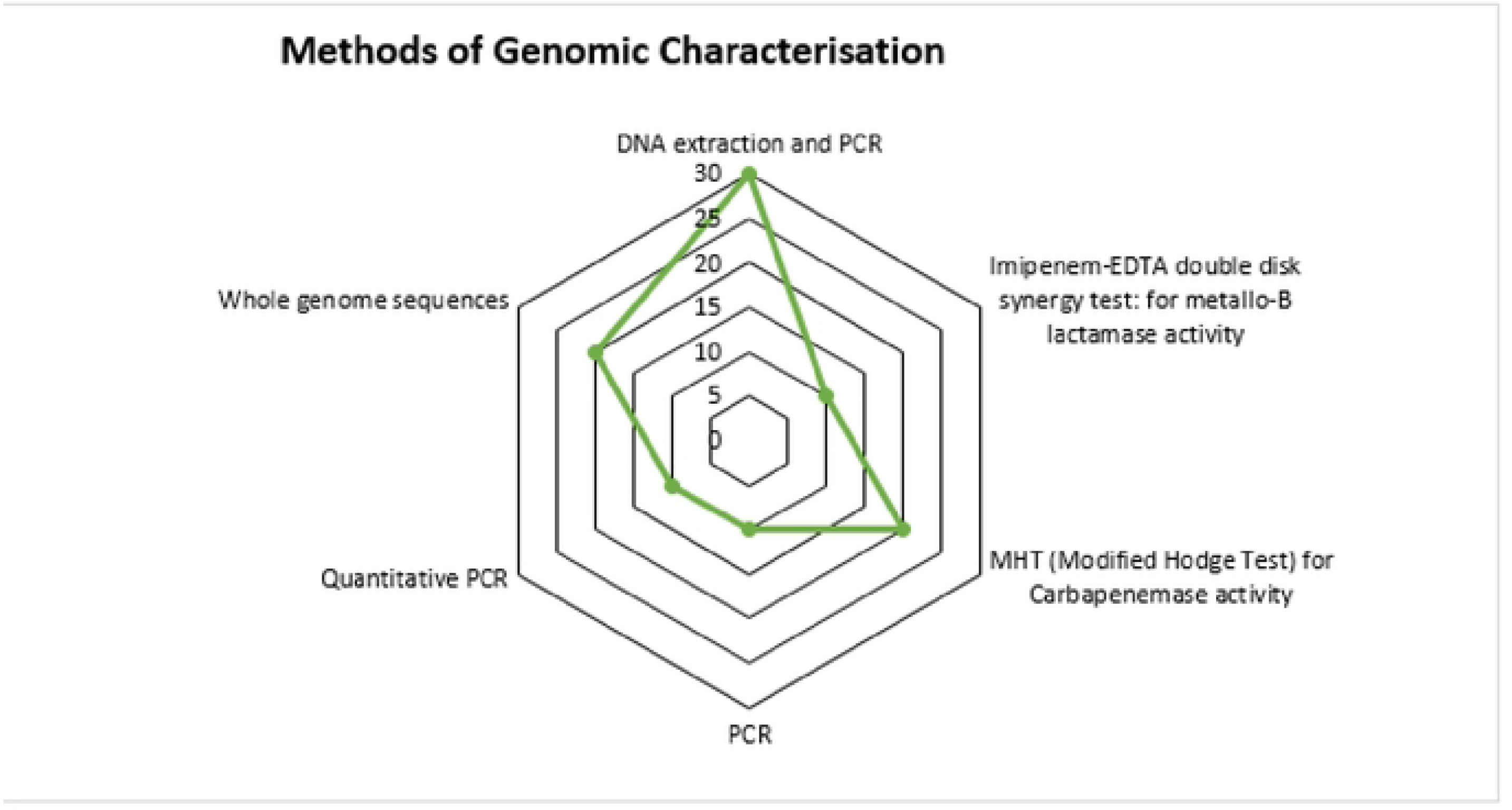
Methods of Genomic Characterization and Sequencing

These antibiotic resistances reported across various studies in East Africa could be driven down by many factors including the unregulated use of antibiotics in the region. Antibiotics are widely available without a prescription in pharmacies and informal marketplaces throughout East Africa. This unrestricted distribution of antibiotics leads to their misuse and overuse, increasing the emergence of resistant strains. Individuals frequently self-medicate, utilize improper doses, or discontinue therapy prematurely, creating an environment favorable to bacterial resistance. The agricultural-wide use of antibiotics in livestock farming in East Africa particularly as growth promoters can also allow resistant bacteria to transfer from animals to humans through the food chain (E.g. milk and meat).

Furthermore, the lack of efficient antimicrobial stewardship programs increases the resistance, as healthcare practitioners frequently use broad-spectrum antibiotics without proper diagnoses. Limited access to second-line drugs, such as amikacin, forces patients to overuse first-line medications, which contributes to resistance. Inadequate infection control procedures and poor cleanliness in healthcare facilities also contribute to the emergence of resistant *Pseudomonas* bacteria. Socioeconomic factors such as poverty and restricted access to healthcare contribute to inappropriate antibiotic usage, while cross-border movement and globalization bring resistant bacteria from other places.

While other antibiotics demonstrated substantial resistance rates, amikacin remained effective in more than 90% of patients (Kiyaga et al., 2022; Moremi et al., 2016). This finding suggests Amikacin’s potential as a treatment for multidrug-resistant *Pseudomonas* infections in the East African Community. Amikacin’s comparatively low resistance may be attributed to its less frequent use compared to other antibiotics in the region. Amikacin, an aminoglycoside, is primarily used for severe infections and is less commonly prescribed than cephalosporins or carbapenems. This limited use helps to sustain its efficacy by reducing the selective pressure for resistance development. Furthermore, because of their potential nephrotoxicity, aminoglycosides need careful dose and monitoring, restricting their normal use and contributing to long-term efficacy.

Furthermore, these findings reveal *Pseudomonas aeruginosa* resistance genes in East Africa Community particularly carbapenem-resistant genes as presented in **Table 3**. The most common carbapenem-resistant genes reported across various studies include *blaNDM-1, blaVIM, blaOXA- 48,* and others, indicating the existence of MBL producers and carbapenemase-resistant bacteria in the region. *VIM* and *NDM* genes, for example, were reported to be present in both clinical isolates and carbapenem-resistant bacteria, indicating the increased threat of carbapenem resistance (Musila et al., 2021; Aruhomukama et al., 2019). These genes produce carbapenemase enzymes, which break down carbapenem antibiotics, making them useless. The growing use of carbapenems as a last resort for serious infections, notably in metropolitan and referral hospitals in the East African Community, has put major selective pressure on *Pseudomonas* populations. This selection pressure, combined with the frequent horizontal gene transfer enabled by mobile genetic elements such as plasmids, has resulted in the fast proliferation of carbapenemase genes. Additionally, the lack of strong infection control procedures in many healthcare settings promotes the transfer of carbapenem-resistant bacteria between patients. This has been compounded by poor wastewater management, particularly in metropolitan areas which allows antibiotic residues and resistant bacteria into the environment, facilitating the spread of resistance genes among environmental microorganisms. This environmental reservoir provides a breeding ground for resistant infections, which can re-enter clinical settings via contaminated water or food.

Additionally, the method utilized to iso++late and identify Pseudomonas spp. has a substantial impact on gene detection for resistance and virulence. Traditional culture was the most common method reported at 70% across various studies in the Community as presented in **Figure 3**. The reason could be due to its low cost and simplicity but it’s limited ability to detect specific genetic resistance mechanisms and virulence factors. Molecular approaches provide a more thorough understanding of resistance and pathogenicity, but they are neglected due to resource and infrastructure constraints. Expanding the use of molecular techniques in East African Community would improve the discovery of important resistance and virulence genes, hence improving patient outcomes and influencing public health efforts to battle *Pseudomonas* infections.

Furthermore, this study found the prevalence of high-risk clones such as ST 244 and ST 357 reported across various studies as presented in **Table 4**. The results demonstrated the clonal spread of multidrug-resistant (MDR) and extensively drug-resistant (XDR) *Pseudomonas aeruginosa* strains across East African Community. These clones include both resistance and virulence genes, making them more harmful in clinical settings (Kiyaga et al., 2022; Musila et al., 2021). The fast clonal proliferation of these high-risk strains is aided by factors such as patient transfers between hospitals, congested healthcare facilities, and inadequate infection control methods. Inadequate sterilization methods and a lack of isolation measures for infected patients allow MDR and XDR clones to spread within and between institutions. Also, the existence of these clones in environmental reservoirs, such as hospital wastewater systems, ensures their persistence and circulation outside clinical settings, which increases the chance of reintroduction into healthcare facilities. The most common method utilized for the detection of these high-risk clones as reported was DNA extraction and PCR at 30%. The reason for this method being increasingly frequent in the East African Community is also+ due to its low cost, availability, simplicity, speed, and effectiveness in detecting specific, therapeutically important resistance genes in resource-limited settings. Other methods reported include quantitative PCR, whole genome sequencing, the Modified Hodge Test (MHT) for carbapenemase activity, the Imipenem-EDTA double disk synergy test for metallo-beta-lactamase activity, and basic PCR as presented in **Figure 5**

*Pseudomonas aeruginosa* reported across various studies in the East African Community had a lower resistance rate (54%) than clinical isolates (73%) Kateete et al. (2016). This finding demonstrates the differences in selecting forces between clinical and environmental settings. This could be because antibiotics are not used extensively in natural habitats, and environmental isolates are subjected to lesser levels of selective pressure, resulting in lower resistance rates. However, environmental bacteria can still serve as repositories for resistance genes, particularly in locations where antibiotic poisoning of water supplies is frequent. As the East African Community faces issues connected to industrial and hospital waste management, the environment becomes a crucial reservoir for resistant strains, which may eventually infect humans through contaminated water or food.

The majority of the studies reporting *Pseudomonas aeruginosa* resistance were conducted in Kenya (30 %) in numerous locations including urban centers like Nairobi and rural areas like Lake Bogoria (Kiyaga et al., 2022). Followed by 30 % in Uganda, next to Kenya in terms of the proportion of studies addressing the topic, indicating a growing awareness of antibiotic resistance and a greater capability for healthcare research. Uganda has made tremendous progress in creating research projects, with institutions like Makerere University and Kampala International University adding to the corpus of knowledge on antibiotic resistance. Studies in Tanzania and the Democratic Republic of the Congo (DRC) are more limited, with an emphasis on specific locations such as Tanzania’s Mwanza both accounting for 20 %. Kenya’s more modern healthcare infrastructure and research institutes offer a more comprehensive examination of resistance patterns in both rural and urban areas. The increased frequency of studies in Kenya may also reflect the country’s dedication to combating antibiotic resistance, which is supported by both local and international research funding. Tanzania and the Democratic Republic of the Congo, on the other hand, confront more obstacles in terms of healthcare resources and research competence, restricting the extent of studies and potentially underestimating the true extent of antimicrobial resistance in these countries as summarized in **Table 5**.

## CONCLUSION

This systematic review revealed the major impact of antibiotics and environment on gene expression patterns of *Pseudomonas spp*. in East Africa Community. The extensive resistance identified, notably in *Pseudomonas aeruginosa* clinical isolates, highlights the importance of unregulated antibiotic use, inadequate healthcare infrastructure, and environmental pollution in generating antimicrobial resistance. Although amikacin is still a potential choice due to its low resistance, the significant incidence of carbapenem-resistant genes (e.g., *blaNDM-1, blaVIM*) and the clonal proliferation of MDR/XDR *Pseudomonas* strains highlight the need for improved infection control and stewardship initiatives. Furthermore, the research finds that, while classic culture methods remain dominant, there is an increasing need to use molecular techniques for improved genomic characterization and resistance identification in the East African Community to improve patient’s outcome.

## RECOMMENDATIONS

1. Healthcare institutions in East Africa must develop strict antimicrobial stewardship programs to monitor and restrict antibiotic consumption, decreasing unnecessary prescriptions and limiting self-medication.
2. Investment in molecular technologies such as quantitative PCR and whole-genome sequencing is required to improve resistance gene and virulence factor detection.
3. To restrict the transmission of MDR/XDR strains, hospitals should establish stronger infection control measures, such as adequate sterilization and isolation protocols.
4. Governments should impose antibiotic-use controls in livestock husbandry to prevent the spread of resistance from animals to people via food.
5. Improve environmental waste management, particularly in metropolitan areas, to prevent the spread of resistant bacteria through water and food systems.

## LIMITATIONS OF THE STUDY

1. While Kenya and Uganda had a large number of studies, data from Tanzania and the DRC were scarce, potentially underestimating overall resistance levels in East Africa.
2. Limited utilization of molecular methods due to resource and infrastructure restrictions may result in underreporting of critical resistance mechanisms.
3. The emphasis on studies published in English or accessible databases may have resulted in a bias that excluded valuable research from other sources.
4. The employment of various antimicrobial sensitivity testing methodologies across studies may have contributed to variations in resistance detection.

